# Angiotensin converting enzyme 2 is a novel target of the γ-secretase complex

**DOI:** 10.1101/2020.09.01.277954

**Authors:** Alberto Bartolomé, Jiani Liang, Pengfei Wang, David D. Ho, Utpal B. Pajvani

## Abstract

Angiotensin converting enzyme 2 (ACE2) is a key regulator of the renin-angiotensin system, but also the functional receptor of the severe acute respiratory syndrome coronavirus 2 (SARS-CoV-2). Based on structural similarity with other γ-secretase (γS) targets, we hypothesized that ACE2 may be affected by γS proteolytic activity. We found that after ectodomain shedding, ACE2 is targeted for intramembrane proteolysis by γS, releasing a soluble ACE2 C-terminal fragment. Consistently, chemical or genetic inhibition of γS results in the accumulation of a membrane-bound fragment of ectodomain-deficient ACE2. Although chemical inhibition of γS does not alter SARS-CoV-2 cell entry, these data point to a novel pathway for cellular ACE2 trafficking.

## Introduction

Angiotensin converting enzyme 2 (ACE2) is a membrane-anchored ectoenzyme that processes Angiotensin II to Angiotensin 1-7, but also mediates the entry of three different coronavirus strains by means of binding the viral spike (S) protein: NL63 (1), SARS-CoV (2) and SARS-CoV-2 (3). S-protein binding to ACE2 triggers membrane fusion and viral entry, but only after S-protein priming by Transmembrane protease serine 2 (TMPRSS2) (3, 4), which also cleaves the ectodomain of ACE2 (5). ACE2 cleavage, or shedding, can additionally be induced by the disintegrin and metallopeptidase domain 17 (ADAM17) (6), which was found to compete with TMPRSS2 (5). In this regard, there are conflicting reports of ADAM17-mediated shedding affecting SARS-CoV entry (5, 7). Viral infection has also been shown to trigger ACE2 endocytosis (8), leading to reduced cell surface expression of ACE2 (9). Intriguingly, ACE2 is seen as a “double-edged sword” (10). While high expression of the receptor enables viral infection, some of the deleterious effects associated with COVID-19 are attributed to loss of ACE2-mediated cardiovascular protection, due to cell surface downregulation (11). In the current COVID-19 pandemic, there has been great interest in novel therapeutics that modulate ACE2, either to prevent SARS-CoV-2 entry (12) or to target the renin-angiotensin system imbalance associated with severe disease (11). Ideally, novel ACE2-focused therapies should be able to disentangle these two faces of the receptor.

The gamma-secretase (γS) protein complex, composed of a Presenilin 1/2 aspartyl protease catalytic core with regulatory (Aph-1a or -1b), enhancer (PEN2) and targeting (Nicastrin) subunits, is the prototype intramembrane-cleaving protease (I-CLiP). I-CLiP proteases introduce a water molecule into the hydrophobic environment of the lipid bilayer for peptide bond hydrolysis within the transmembrane domain. γS targets are typically single-pass, type I transmembrane proteins with large ectodomains and C-terminal intracellular domains (ICD). γS substrates first undergo ectodomain shedding at the cell surface, rendering a membrane-bound protein stub that is targeted by γS for intramembrane proteolysis. The released soluble ICD tends to be rapidly degraded by the proteasome, but in some cases, such as the Notch family of cell surface receptors (13) and the amyloid β precursor protein (APP) (14), the ICD has signaling activity. For example, Notch ICD binds a Mastermind/Rbpj complex to activate transcription of canonical Notch target genes (15). But in addition to Notch and APP, dozens of other putative γS targets have been identified (16), not determined by an amino acid consensus sequence, but rather specific transmembrane conformational structure and accessibility (17, 18). Validation of novel γS targets is hindered by the lack of clear-cut common features and ectodomain shedding requirements.

Based on structural similarity of ACE2 to known γS targets, we hypothesized that γS regulates intramembrane cleavage of ACE2 and may impact SARS-CoV-2 biology. Here we report that ACE2 undergoes TMPRSS2/ADAM17-dependent γS cleavage, resulting in a short-lived ACE2-ICD. Genetic or chemical inhibition of γS prevents ACE2-ICD generation, leading to accumulation of a membrane-bound ACE2 lacking the ectodomain. However, we show using a pseudovirus system that γS inhibition does not impact SARS-CoV-2 cellular entry.

### Experimental Procedures

#### Antibodies and chemicals

We used antibodies against GFP (B-2) sc-9996, Nicastrin (N-19) sc-14369, Rhodopsin (ID4) sc-57432, C9 tag (TETSQVAPA peptide) from Santa Cruz Biotechnology; Actin, A2066 from Millipore-Sigma; Presenilin 1 (D39D1) 5643, from Cell Signaling Technology; and ACE2 ab15348, from Abcam. We used MG132 and phorbol 12-myristate 13-acetate (PMA) (Sigma). γS inhibitors (GSI) were Compound E (Axxora) and dibenzazepine (DBZ) (Syncom). DBZ was used in all experiments indicating GSI usage unless otherwise stated.

#### Cell culture and cell lines

We used Presenilin-deficient (Psen1/2 double knockout) and control mouse embryonic fibroblasts (MEFs) provided by Nikolaos Robakis (Mount Sinai School of Medicine, New York, NY) (19) and Nicastrin knockout and control MEFs were obtained from Phillip Wong (Johns Hopkins University School of Medicine, Baltimore, MD) (20). We cultured MEFs, Caco-2, VeroE6 and 293T cells in DMEM supplemented with 10% heat-inactivated fetal bovine serum (FBS) and 1% penicillin-streptomycin (Thermo-Fisher). For transfection experiments, we used Lipofectamine 3000 and OptiMEM (Thermo-Fisher) as per the manufacturer’s instructions.

#### Plasmids

We obtained a vector encoding C-terminally tagged ACE2 (TETSQVAPA, C9-tag) from Hyeryun Choe, obtained from Addgene (#1786) (2). We replaced C9 with EGFP to generate ACE2-GFP, which we in turn deposited on Addgene (#154962). We obtained a TMPRSS2 expression vector from Roger Reeves, obtained from Addgene (21).

#### Western blotting, immunoprecipitation and quantitative PCR

We lysed cells in RIPA buffer containing protease inhibitors (Pierce protease inhibitor tablets, Thermo-Fisher), and 10 mM NaF. For immunoprecipitation of γS, we lysed cells in 1% CHAPSO, 100 mM NaCl, 2 mM EDTA, 25 mM Tris-HCl (pH 7.4), with protease inhibitors, and immunoprecipitated 1.2 mg of protein lysate with 2.5 μg C9-tag antibody and Protein G magnetic beads (Cell Signaling Biotechnology). After overnight incubation, we separated beads with a DynaMag-2 magnet (Thermo-Fisher), and washed three times in buffer containing 0.5% CHAPSO. We resuspended beads in 2× Laemmli buffer and heated at 70°C for 10 min, prior to SDS-PAGE, Western blot and visualization with the ECL Western Blotting Detection Kit (GE Healthcare Bio-Sciences). We performed qPCR as previously described (22) with primers specific for human (h) Caco-2 cells; or *Chlorocebus sabaeus* (cs) VeroE6 cells as follows: *h/csACE2:* TGGTGGGAGATGAAGCGAGA, AACATGGAACAGAGATGCGGG; *hTMPRSS2:* CACCGAGGAGAAAGGGAAGAC, CATGGCTGGTGTGATCAGGT; *csADAM17*: AGGTGTCCAGTGCAGTGATAGG, ATCTTCAGCATTTCCCGGAGG; *hADAM17:* CGTTGGGTCTGTCCTGGTTT, TCAGCATTTCGACGTTACTGGG. We normalized with peptidylprolyl isomerase A using the following primers: *csPPIA:* CAGGTCCTGGCATCTTGTCC, GCTTGCCATCCAACCACTCA; *hPPIA:* TATCTGCACTGCCAAGACTGAGTG, CTTCTTGCTGGTCTTGCCATTCC.

#### Immunofluorescence and confocal imaging

We seeded cells on glass coverslips as previously described (23), and gathered images with an Axio Observer Z1 with an LSM 710 scanning module (Zeiss), collected using a 63× Zeiss Plan-Apochromat oil objective. All images were obtained in a 1,024-by 1,024-pixel format and processed with ZEN2 (Zeiss).

#### SARS-CoV-2 pseudovirus and cell entry inhibition

Recombinant Indiana vesicular stomatitis virus (rVSV) expressing SARS-CoV-2 S-protein, and the neutralizing antibody used as control, were generated as previously described (24). We grew 293T cells to 80% confluency before transfection with pCMV3-SARS-CoV-2-spike using FuGENE 6 (Promega), and cultured cells overnight at 37°C with 5% CO_2_. The next day, we removed medium and infected cells with VSV-G pseudo-typed ΔG-luciferase (G*ΔG-luciferase, Kerafast) in DMEM at an MOI of 3 for 1 hour before washing the cells with 1×DPBS three times. We added DMEM supplemented with anti-VSV-G antibody (I1, mouse hybridoma supernatant from CRL-2700; ATCC) to the infected cells and harvested the supernatant the next day. To test DBZ inhibition of SARS-CoV-2 cell entry, we seeded VeroE6 or Caco-2 cells in a 96-well plate at a concentration of 2 × 10^4^ cells per well. We incubated pseudovirus the next day with serial dilutions of DBZ in triplicate for 30 min at 37 °C. We added the mixture to cultured cells and incubated for an additional 24 h, with an S-protein neutralizing antibody was used as control (24). We measured luminescence using a Britelite plus Reporter Gene Assay System (PerkinElmer).

## RESULTS

### ACE2 ectodomain shedding is required for γS cleavage

Consistent with other confirmed γS targets (i.e. APP (14), Notch (13) and Jagged1 (25)), ACE2 has a large ectodomain that can be processed by a sheddase (ADAM17/TMPRSS2) (5, 6) and a single transmembrane domain (Fig. 1A). Based on this structural similarity, we hypothesized that after ectodomain shedding, the resultant protein (ACE2ΔE) may represent a novel γS target. To test this hypothesis, we expressed ACE2 tagged at its C-terminus in 293T cells, and triggered ectodomain shedding by either TMPRSS2 co-expression, or PMA-induced activation of endogenous sheddases (26). In the presence of TMPRSS2, we observed a 15 kDa C-terminal ACE2 fragment which accumulated in the presence of a γS inhibitor (GSI), dibenzazepine (DBZ) (Fig. 1B). PMA treatment rendered a 18 kDa C-terminal fragment, consistent with the expected size after ADAM17-mediated ACE2 cleavage (6). Similar to TMPRSS2 co-expression, this longer form of ACE2ΔE also accumulated in DBZ treated cells (Fig. 1C). Intriguingly, we observed ACE2ΔE accumulation even in unstimulated cells expressing ACE2 (Fig. 1B, C), suggesting endogenous ectodomain shedding followed by γS cleavage is part of the normal turnover of ACE2. We also used a chemically distinct GSI (Compound E) (Fig. 1D), that confirmed that ACE2ΔE is targeted by γS.

**Figure 1.**
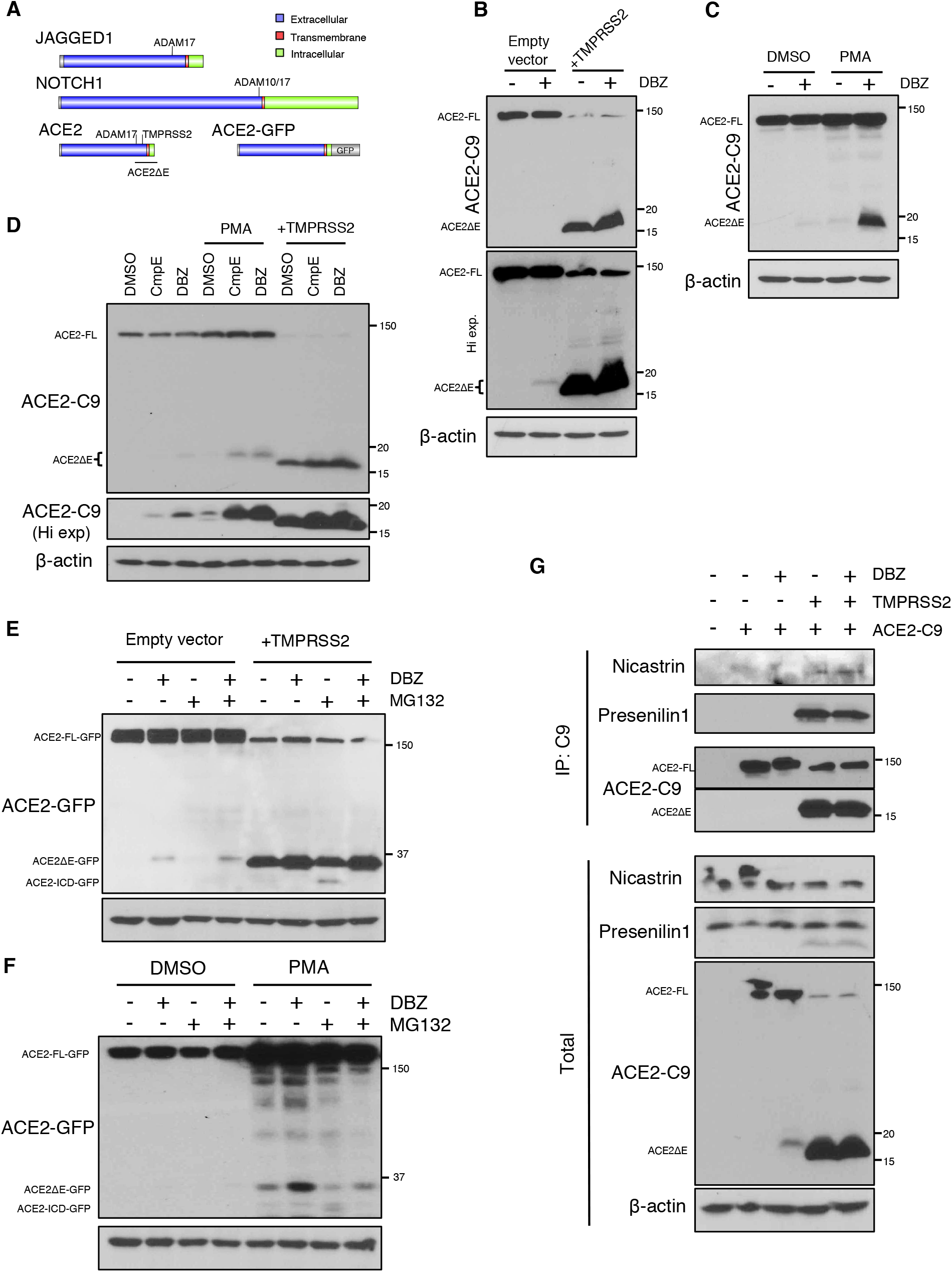
ACE2 is targeted by γS after ectodomain shedding. A, Schematic and scaled representation of γS targets JAGGED1 or NOTCH1, with ACE2 and ACE2-GFP. Domains and regions targeted by sheddases are depicted. B, Western blots from 293T cells transfected with ACE2-C9 with or without TMPRSS2, then treated with DBZ (dibenzazepine 100 nM) (+), or DMSO (-). Mobility consistent with full length (FL) and ACE2 lacking its ectodomain (ACE2ΔE) indicated. C, Western blots from 293T cells transfected with ACE2-C9 with or without PMA (200 nM, 15h) treatment. D, Western blots from 293T cells transfected with ACE-C9, with or without TMPRSS2, or treated with PMA, with or without two γS inhibitors [DBZ (dibenzazepine, 100 nM) or CmpE (compound E, 40 nM)]. E, Western blots from 293T cells transfected with ACE2-GFP, with or without TMPRSS2, then treated with DBZ and/or MG132 (1 µM, 15h) F, Western blots from 293T cells transfected with ACE2-GFP, with or without PMA, DBZ and/or MG132. G, Western blots from immunoprecipitates derived from 293T cells transfected with ACE2-C9, with or without TMPRSS2, then treated with DBZ. Data is representative of 2-3 independent experiments.

Most γS-liberated target protein ICDs are extremely labile and rapidly degraded in the proteasome (16). As the cytoplasmic portion of ACE2 is too small for conventional SDS-PAGE, we generated a C-terminal ACE2-GFP fusion protein to detect ACE2-ICD production. We repeated the above experiments using this novel construct and found that TMPRS2 co-expression or PMA treatment provoked ectodomain shedding and ACE2ΔE-GFP accumulation with DBZ treatment (Fig. 1E, F). As hypothesized, we also observed ACE2-ICD-GFP only after proteasome inhibition (Fig 1E, F).

Based on these pharmacologic data, we next hypothesized that ACE2ΔE and γS would physically interact. To test this, we performed co-immunoprecipitation of endogenous γS with C-terminally tagged ACE2, and observed association with both Nicastrin and Presenilin1 with ACE2ΔE but not full-length ACE2, consistent with other *bona fide* γS targets (18). γS-ACE2ΔE interaction did not change in the presence of DBZ (Fig. 1G). In sum, these data establish ACE2 as a novel γS target.

### ACE2 cleavage-dependent localization is altered in γS-deficient cells

To determine cellular ramifications of γS-mediated ACE2 cleavage, we next evaluated ACE2 processing in Nicastrin knockout (Ncstn KO) (20) or Presenilin1/2 double knockout (Psen1/2 dKO) MEFs (19), both of which have disrupted γS activity. Consistent with GSI treatment, these γS-deficient cell lines displayed ACE2ΔE accumulation, accentuated by co-expression of TMPRSS2 (Fig. 2A). In this experimental paradigm, the C-terminus of ACE2 is primarily localized to the membrane but appeared diffusely cytoplasmic with TMPRSS2 expression in control cells (Fig. 2B and 2C, top panels). In γS-deficient cells however, ACE2 remained membrane-associated even in the presence of TMPRSS2 (Fig. 2B and 2C, bottom panels). We reproduced these results using a C-terminal ACE2-GFP fusion protein (Fig. 2D), and with DBZ treatment of γS in control cells (Fig. 2E). These results indicate that γS is required for the release of a soluble C-terminal ACE2 fragment from cell membranes.

**Figure 2.**
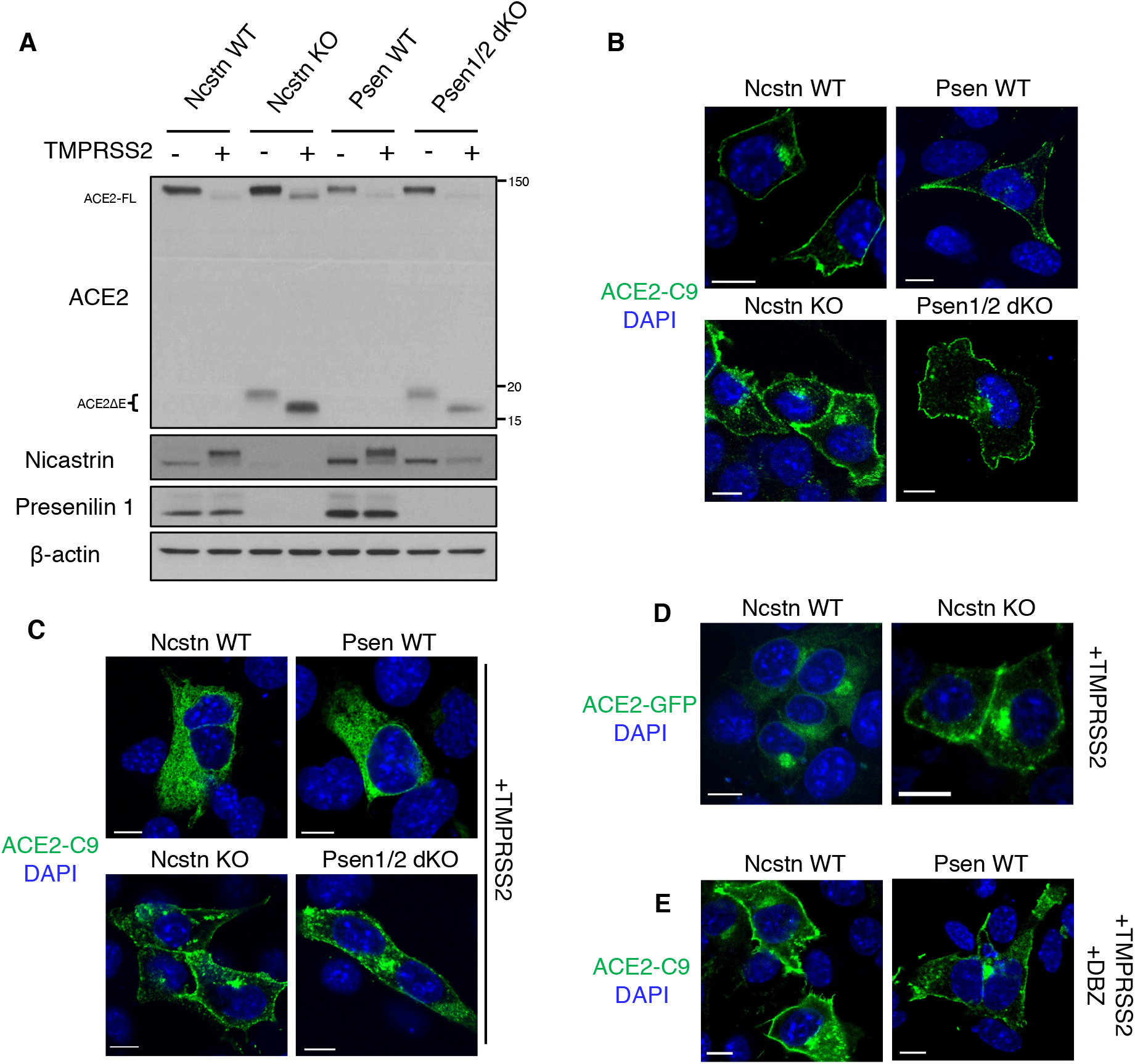
γS-deficient cells cannot process ACE2ΔE. A, Western blots from Presenilin 1/2 double KO (Psen1/2 dKO) or Nicastrin KO (Ncstn KO) MEFs and their wild type (WT) controls. Cells were transfected with ACE2-C9 with or without TMPRSS2. B, Representative immunofluorescence images of WT MEFs transfected with ACE2-C9. C, Representative immunofluorescence images of WT, Psen1/2 dKO and Ncstrn KO MEFs transfected with ACE2-C9 and TMPRSS2. D, GFP fluorescence in WT or Ncstrn KO MEFs co-expressing ACE2-GFP and TMPRSS2. E, Representative immunofluorescence images of WT MEFs transfected with ACE2-C9 and TMPRSS2 in the presence of GSI. Scale bars: 10 µm.

### Endogenous ACE2 cleavage is regulated by γS

293T and MEFs do not express significant levels of endogenous ACE2. To confirm the physiologic relevance of γS-mediated ACE2 cleavage, we used two well-characterized ACE2-positive cell lines that allow SARS-CoV-2 infection and replication, Caco-2 and VeroE6. Using an antibody that recognizes the C-terminal region of ACE2, we observed accumulation of endogenous ACE2ΔE with GSI treatment in both cell lines (Fig. 3A-C). These data confirmed results from ectopic ACE2 expression. We next took advantage of this system to test whether longer ACE2ΔE half-life or impaired production of ACE2-ICD may produce negative feedback on this pathway in these cells. This hypothesis was based on nuclear localization and transcriptional activity of the C-terminal fragment of the related protein, ACE (27, 28).

**Figure 3.**
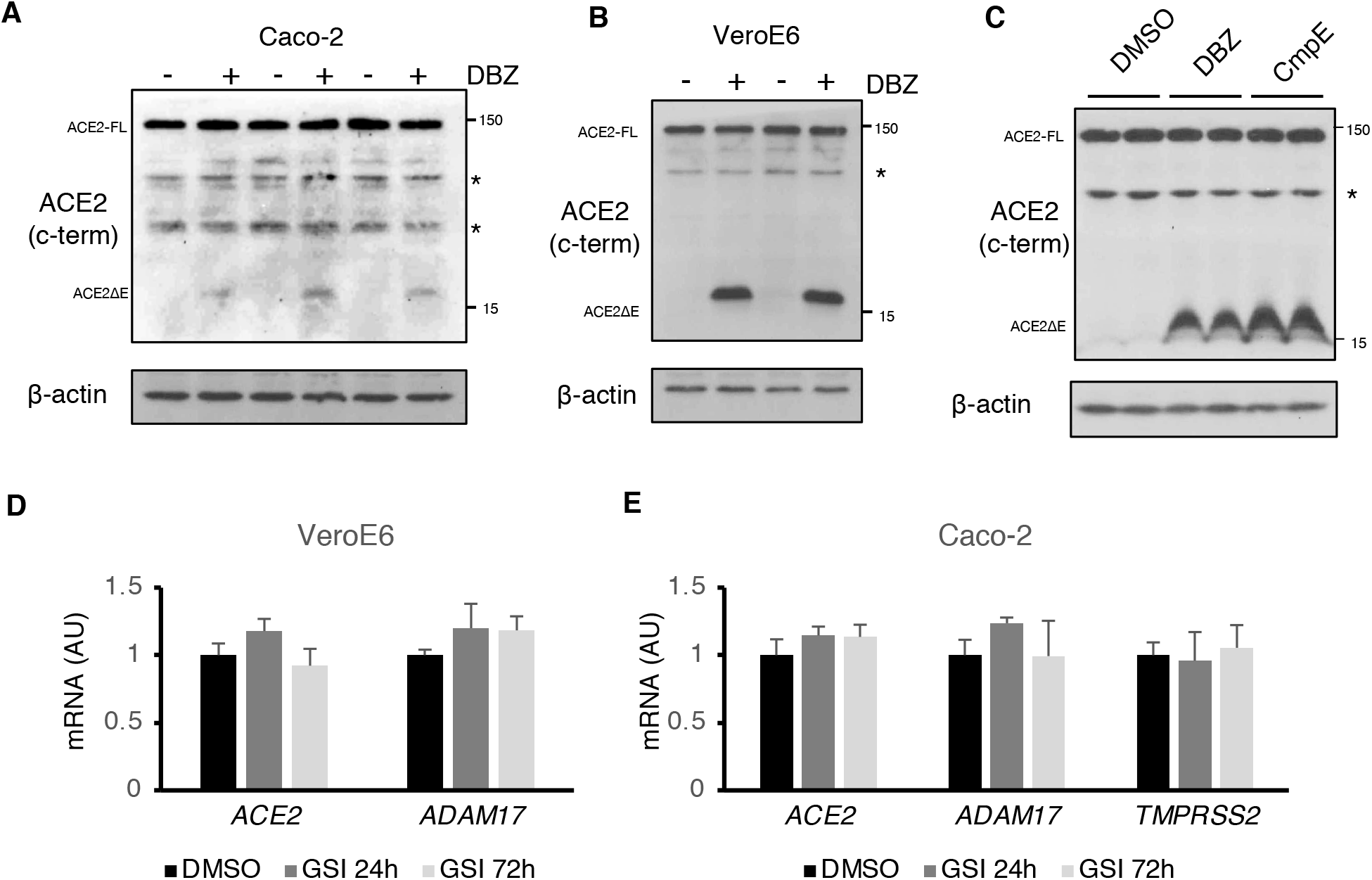
γS inhibition targets endogenous ACE2. A, Western blots of Caco-2 cells treated with DBZ (+) or DMSO (-). B, Western blots of VeroE6 cells treated with DBZ (+) or DMSO (-). C, Western blots of VeroE6 cells treated with DBZ compound E (CmpE) or DMSO (-). D, Gene expression in VeroE6after 24 or 72h treatment with DBZ, showing means ± SD. E, Gene expression in Caco-2 cells after 24 or 72h treatment with DBZ, showing means ± SD.

However, we did not observe nuclear ACE2 or differences in expression of *ACE2, TMPRSS2* or *ADAM17* in DBZ-treated VeroE6 or Caco-2 cells (Fig. 3D-F). These data render unlikely the possibility that ACE2-ICD mediates feedback inhibition on *ACE2* gene expression.

### γS inhibition does not alter SARS-CoV-2 S-protein-mediated cell entry

As genetic or pharmacologic γS inhibition affected ACE2 cleavage and subcellular localization, we hypothesized that GSI may reduce SARS-CoV-2 cell entry and replication. To test this potential, we utilized SARS-CoV-2 S-protein pseudotyped with VSV and tested a wide range of DBZ concentrations (0.3 nM-1 µM). In comparison to a potent S-protein neutralizing antibody, used as a positive control (24), DBZ did not affect SARS-CoV-2 S-protein mediated viral entry in VeroE6 or in Caco-2 cells (Fig. 4). These results indicate that although γS is necessary for ACE2 intracellular processing, blocking γS does not affect viral entry.

**Figure 4.**
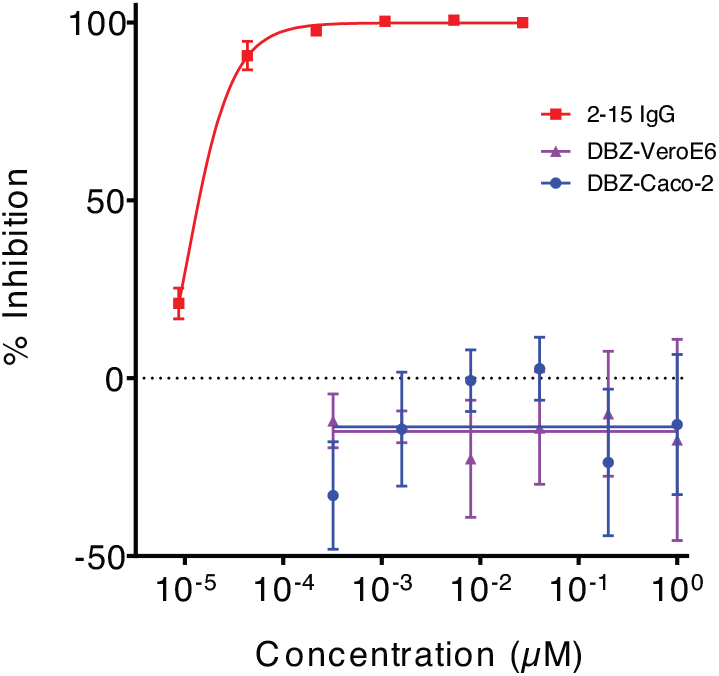
γS inhibition does not prevent SARS-CoV-2 S-protein mediated cell entry. Inhibition of SARS-CoV-2 pseudovirus by DBZ at the indicated concentrations tested on VeroE6 or Caco-2 cells. A SARS-CoV-2 neutralizing antibody, 2-15, tested on VeroE6 was used as a positive control. Triplicates are presented as means ± SEM.

## Discussion

ACE2 has recently caught the attention of the research community because of its role as the functional receptor of SARS-CoV-2 (3). Here we have characterized ACE2 as a novel target of γS (Fig. 5). Similar to other known targets (16-18), ectodomain shedding prompts γS-mediated intramembrane cleavage to release soluble ACE2-ICD. Some ICDs (i.e. Notch) generated by γS are transcriptionally active, but a functional role of many others remains elusive (16). γS has also been dubbed as the “proteasome of the membrane” (29). Our finding that ACE2-ICD is rapidly cleared by proteasomal degradation suggests is consistent with the view that γS-mediated cleavage represents a way to dispose membrane proteins stubs. However, our data cannot as yet discard the hypothesis that ACE2-ICD might represent a novel biologically active peptide.

**Figure 5.**
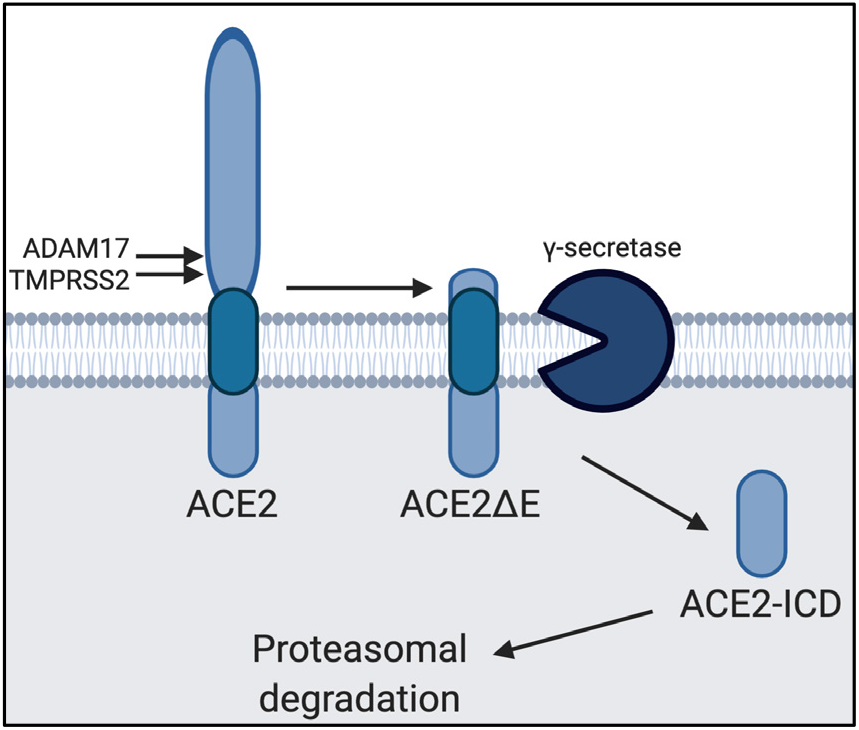
Model of ACE2 cleavage. Model showing the sequential processing of full length ACE2 by ADAM17/TMPRSS2 and γS, rendering ACE2ΔE and ACE2-ICD, respectively. ACE2-ICD is then rapidly degraded in the proteasome.

Previous reports indicate that ACE2 processing by cell membrane proteases such as ADAM17 or TMPRSS2 impacts SARS-CoV S-protein mediated cell entry (5, 7). Our data is clear that pharmacologic inhibition of γS-mediated ACE2 cleavage does not, but cannot rule out the possibility that other compounds termed “γS modulators” (GSMs) (30) may behave differently. GSMs, developed primarily to differentially affect γS processing of so-called “on-target” (i.e. APP) as opposed to “off-target” (i.e. Notch) substrates, may in fact selectively increase γS processivity. In light of our finding that ACE2 is a novel γS target, GSMs are worth evaluating for biological activity against SARS-CoV-2 pathogenesis. In addition, some groups have speculated that blocking Notch signaling with GSIs may ameliorate COVID-19 progression (31). Notch signaling promotes M1 polarization of macrophages (32), and also contributes to T-cell cytokine production (33). Thus, despite our finding that GSI does not directly affect viral entry, potential effects to block Notch-induced hyperinflammation suggest compounds that have completed Ph2/3 clinical trials can potentially be repurposed for COVID-19.

In addition to the relatively recently discovered role as a viral receptor, ACE2 has known roles in the renin-angiotensin system (34), but also other potential functions. For example, mutations causal of Hartnup disorder impair association of the neutral amino acid transporter SLC6A19 with ACE2, suggesting that ACE2 serves as a chaperone for membrane trafficking (35), akin to the function of collectrin towards SLC6A19 or SLC1A1 (36). It is possible that γS-mediated transmembrane processing of ACE2 may impact ACE2 chaperone ability, or even in the structurally homologous collectrin. These potential ramifications of our findings require further study.

In sum, our results demonstrate that ACE2 is a novel γS target, but that pharmacologic inhibition of γS does not impact SARS-CoV-2 S-protein mediated cell entry. Given the pharmacologic accessibility of γS, with prior evaluation of GSIs and GSMs for Alzheimer’s Disease and cancer, we present these data to encourage further exploration into this novel biology for application to COVID-19 or to other pathology attributable to the myriad functions ascribed to ACE2.

## Acknowledgments

We thank Michael Yin, and members of the Pajvani and Ho laboratories for insightful discussion.

## Funding and additional information

Supported by NIH DK103818 (UBP) and a Russell Berrie Foundation Fellowship in Diabetes Research (AB). The content is solely the responsibility of the authors and does not necessarily represent the official views of the National Institutes of Health.

## Author contributions

A.B. designed, performed and interpreted experiments, and wrote the manuscript. J.L., P.W., and D.D.H performed and interpreted experiments. U.B.P. designed and interpreted experiments, and wrote the manuscript.

## Data and materials availability

All data associated with this study are present in the manuscript.

